# Autonomic and neurotrophic signaling in alcohol-associated liver disease

**DOI:** 10.64898/2026.09.27.754815

**Authors:** Yingjie Zhang, Jiaqi Liu, Furong Zhang, Montufar-Solis Dina, Yukun Guan, Ting Wu, Yuichiro Mori, Sean Marrelli, Guangwei Du, Caitlin Thornley, Bin Gao, Mark Hobeika, Xiaoqin Wu

**Author notes:** **Corresponding author**: Xiaoqin Wu, PhD, Department of Integrative Biology and Pharmacology McGovern Medical School, UTHealth at Houston 6431 Fannin Street MSB4.212, Houston, TX 77030 Email: mailto.

## Abstract

Alcohol-associated liver disease (ALD) is a leading cause of liver-related morbidity and mortality, yet its pathogenesis remains incompletely understood. While neural regulation is critical for hepatic homeostasis, how autonomic and neurotrophic signaling is regulated in the liver during ALD progression remains largely unexplored. Here, we performed an integrative analysis of bulk and single-cell transcriptomic datasets of human livers across the spectrum of ALD. We identified stage-dependent remodeling and coordinated expression patterns of autonomic and neurotrophic genes. Autonomic signaling exhibited a consistent decline, with reduced expression of sympathetic (*DBH*, *TH*) and the parasympathetic (*ChAT*) markers. In contrast, pan-neuronal and neuroplasticity-associated genes (*UCHL1*, *TUBB3*, *STMN2*) were upregulated, indicating active neural remodeling. In parallel, factors such as *MANF*, *GDNF*, and *NRG2*, with the receptor *NTRK1*, were upregulated and positively correlated with clinical indicators of disease severity, inflammation and fibrosis. In contrast, trophic factors including *NTF3* and *CDNF*, and receptors such as *GFRA1* and *GFRA3*, were downregulated and inversely associated with these parameters. Single-cell analyses revealed that neurotrophic factors are predominantly expressed by hepatic stellate cells, supporting their role as resident neuroglia-like cells in the liver. In another independent cohort, we observed a coordinated downregulation of NTF3 and sympathetic nerve markers in alcohol-associated hepatitis (AH), suggesting impaired NTF3-mediated neurotrophic support may contribute to sympathetic dysfunction in AH. Collectively, these findings uncover previously unrecognized neuro-regulatory dimensions in ALD and suggest that dysregulated autonomic and neurotrophic signaling may contribute to disease progression.

**Conflict of interest:** The authors have declared that no conflict of interest exists.

## Introduction

Alcohol-associated liver disease (ALD) represents a major and growing global health burden and remains a leading cause for liver transplantation and liver-related mortality worldwide. Recent epidemiological analyses indicate that approximately 5.1% of adults worldwide have alcohol use disorder (AUD). ALD now accounts for an estimated 354,250 deaths, representing a 32.6% increase since 2000, and an additional 92,230 deaths from alcohol-attributable primary liver cancer (1, 2). ALD encompasses a spectrum from simple steatosis to alcohol-alcohol-associated hepatitis (AH), fibrosis, cirrhosis, and hepatocellular carcinoma. Despite significant advances in defining molecular and inflammatory pathways involved in ALD, effective therapeutic options for severe AH remain limited, indicating the incomplete understanding of disease pathogenesis.

Beyond its direct hepatotoxic effects, chronic alcohol exposure exerts profound systemic effects through autonomic nervous system (3). In peripheral tissues, alcohol consumption increases muscle sympathetic nerve activity, contributing to cardiovascular dysfunction (4). In metabolic organs, binge or chronic ethanol exposure activates norepinephrine–β3-adrenergic receptor (Adrb3) signaling in adipocytes, promoting lipolysis and elevating circulating free fatty acids, which in turn exacerbate ethanol-induced liver injury (5, 6). Despite these well-established neuro-metabolic interactions in other organ systems, the direct impact of alcohol exposure on hepatic neural networks, particularly sympathetic and parasympathetic pathways, remains poorly understood. The liver is a highly innervated organ that integrates autonomic and sensory neural inputs to regulate metabolic homeostasis, immune responses, and tissue repair following injury. Recent advances in three-dimensional imaging have revealed that sympathetic innervation predominates in both mice and human livers, whereas parasympathetic inputs are sparse or absent under physiological conditions (7, 8). Neurotrophic factors, classically recognized for their roles in neuronal development, survival, and plasticity (9), have emerged as critical regulators of cellular stress responses and tissue homeostasis in peripheral organs, including the liver (10–12). While alcohol-induced alterations in neurotrophic signaling are well documented in the central nervous system, how these pathways are regulated in the liver during ALD progression and they interact with hepatic autonomic circuits remain largely unexplored.

To address this knowledge gap, we performed an integrative transcriptomic analysis of hepatic autonomic pathways and neurotrophic signaling using bulk and single-cell RNA-sequencing (scRNA-seq) datasets from healthy controls (HCs) and patients across distinct stages of ALD and cirrhosis. Specifically, we sought to address three key questions: (i) whether autonomic nerve pathways and neurotrophic signaling exhibits disease–, stage– and cell-specific transcriptional signatures during ALD progression; (ii) whether these signatures are intrinsic correlated and associated with hepatic inflammation and injury; and (iii) whether they reflect or potentially mediate autonomic imbalance characteristics of ALD. Our findings reveal distinct transcriptional programs of autonomic nervous system and neurotrophic signaling that strongly correlate with clinical severity and inflammatory pathways in human ALD, highlighting their potential roles as previously underappreciated modulators of disease progression. Importantly, these transcription patterns were validated at the protein level in an independent cohort of human liver samples, supporting their robustness and translational relevance. Collectively, this work provides a conceptual framework linking autonomic nerve system and neurotrophic signaling to hepatic pathophysiology in ALD.

## Results

### Autonomic nerve markers are progressively dysregulated during ALD progression and associated with clinical severity

We first investigated whether hepatic nerve-specific gene signatures are altered in response to chronic alcohol exposure and during ALD progression. Strikingly, we observed that canonical sympathetic nerve-associated genes, including dopamine β-hydroxylase (*DBH*), tyrosine hydroxylase (*TH*), and solute carrier family 6 member 2 (*SLC6A2*), were significantly downregulated in patients with sAH and ExAH. In contrast, pan-neuronal and neuroplasticity-associated markers, such as tubulin beta-3 class III (*TUBB3*), stathmin-2 (*STMN2*), and ubiquitin C-terminal hydrolase L1 (*UCHL1*), exhibited a progressive upregulation across ALD stages (**Figure 1A&1B**).

**Figure 1.**
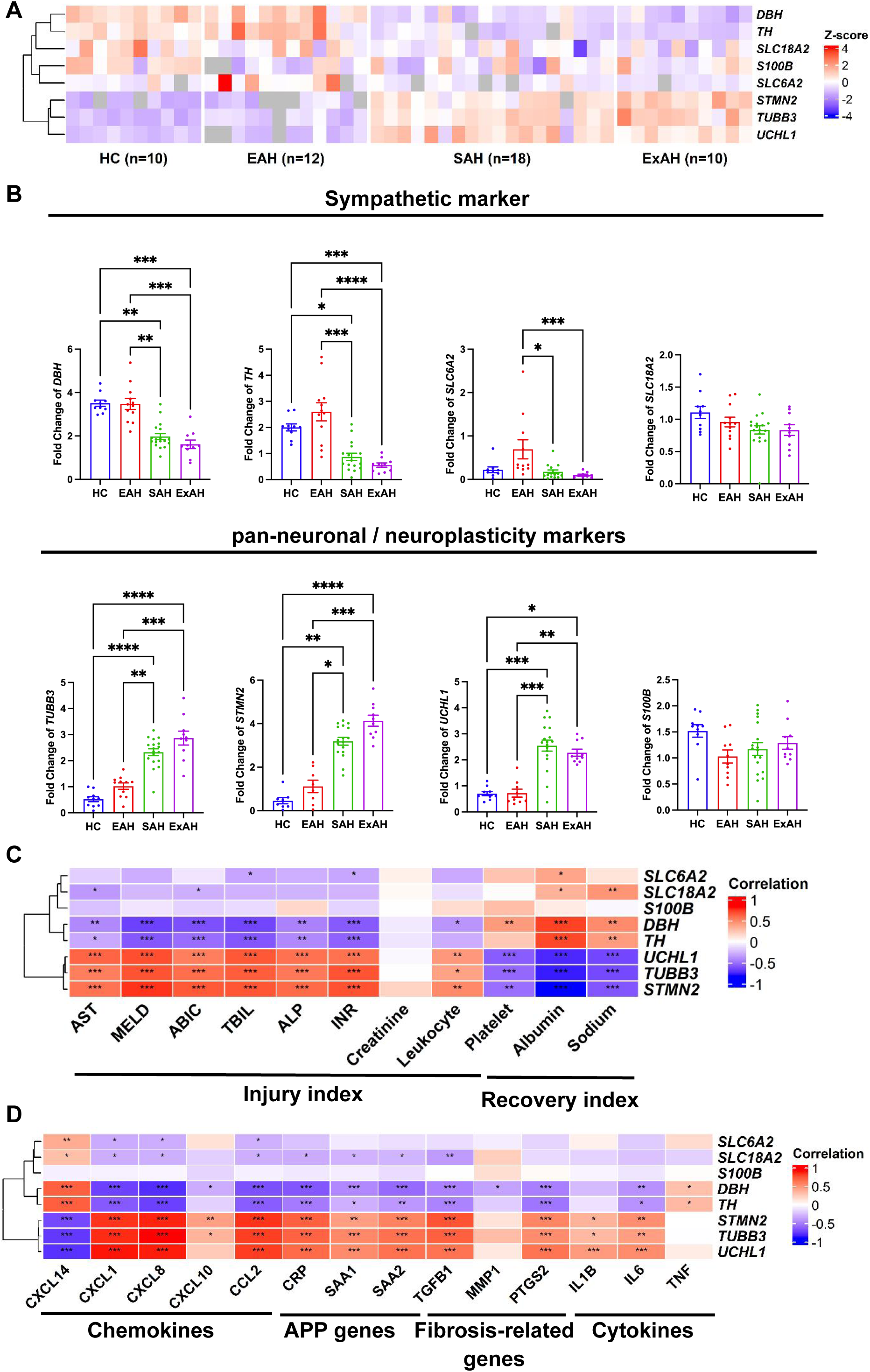
Neural marker expression and their associations with clinical parameters in patients with ALD. (**A**) Heatmap showing Z-score–normalized expression of neural markers across ALD stages (HC, EAH, SAH, ExAH). (**B**) Bar plots showing the expression levels of sympathetic and pan-neuronal/neuroplasticity markers across different groups. Each dot represents an individual sample. (**C**) Heatmap showing correlations between neural markers and clinical parameters, including liver injury, disease severity, and recovery indices. (**D**) Heatmap showing correlations between neural markers and inflammation– and fibrosis-related gene signatures. Correlations were calculated using Spearman’s rank correlation. Statistical analysis for group comparisons was performed using the Kruskal–Wallis test followed by Dunn’s multiple-comparison test. *\*P < 0.05, **P < 0.01, ***P < 0.001, ****P < 0.0001*.

To further define the clinical relevance of these changes, we performed correlation analyses with key indicators of liver injury and inflammation. Sympathetic neural markers showed strong negative correlations with serum markers of hepatic injury, pro-inflammatory cytokines and chemokines, as well as fibrosis-associated genes (**Figure 1C&1D**). In contrast, pan-neuronal and neuroplasticity markers were positively associated with these pathological parameters (**Figure 1C&1D**). Notably, *DBH* and *TH* showed positive correlations with cytokine *TNF-α* levels (**Figure 1D**). Together, these findings reveal a coordinated yet divergent remodeling of hepatic neural signatures during ALD progression, characterized by suppression of sympathetic neural identity alongside increased expression of pan-neuronal and injury-associated neuroplasticity markers. This shift may reflect both loss of functional sympathetic innervation and activation of stress– or injury-induced neural remodeling programs and is closely linked to disease severity and hepatic inflammation.

### Distinct transcriptional signatures of neurotrophic factors in human ALD

Neurotrophic factors play essential roles in neuronal survival, growth, differentiation, and functional maintenance (13). To systemically characterize the transcriptomic profiles of neurotrophic signaling across the different stages of ALD, we analyzed a publicly available liver bulk RNA-seq dataset, which includes human samples from healthy controls (HC), early alcoholic hepatitis (EAH), severe alcoholic hepatitis (SAH), and explanted end-stage alcoholic livers (ExAH). Principal component analysis (PCA) based on a predefined panel of 26 neurotrophic factors revealed clear separation among ALD disease stages (**Figure 2A**), indicating that stage-specific neurotrophic factor expression patterns are closely associated with ALD progression. Consistently, Z-score-normalized heatmap analysis of neurotrophic factors and their receptors demonstrated distinct expression signatures across disease stages (**Figure 2B**). Notably, multiple neurotrophic factors including ciliary neurotrophic family (*CNTF*), mesencephalic astrocyte-derived neurotrophic factor (*MANF*), neuregulin-2 (*NRG2*) were progressively upregulated as disease advanced, whereas others, such as nerve growth factor (*NGF*), neurotrophin-3 (*NTF3*) and cerebral dopamine neurotrophic factor (*CDNF*) showed gradual downregulation during ALD progression (**Figure 2C**). In addition, the transcriptional levels of neurotrophic factor receptors exhibited stage-dependent alterations (**Figure 2D**). For example, most receptors, except neurotrophic receptor tyrosine kinase 1 (*NTRK1*), were downregulated as the disease became more severe, while rearranged during transfection (*RET)* expression was transiently induced in EAH, but returned to baseline in SAH and ExAH. Together, these findings indicate that neurotrophic signaling undergoes dynamic stage-specific transcriptional reprogramming during the progression of ALD.

**Figure 2.**
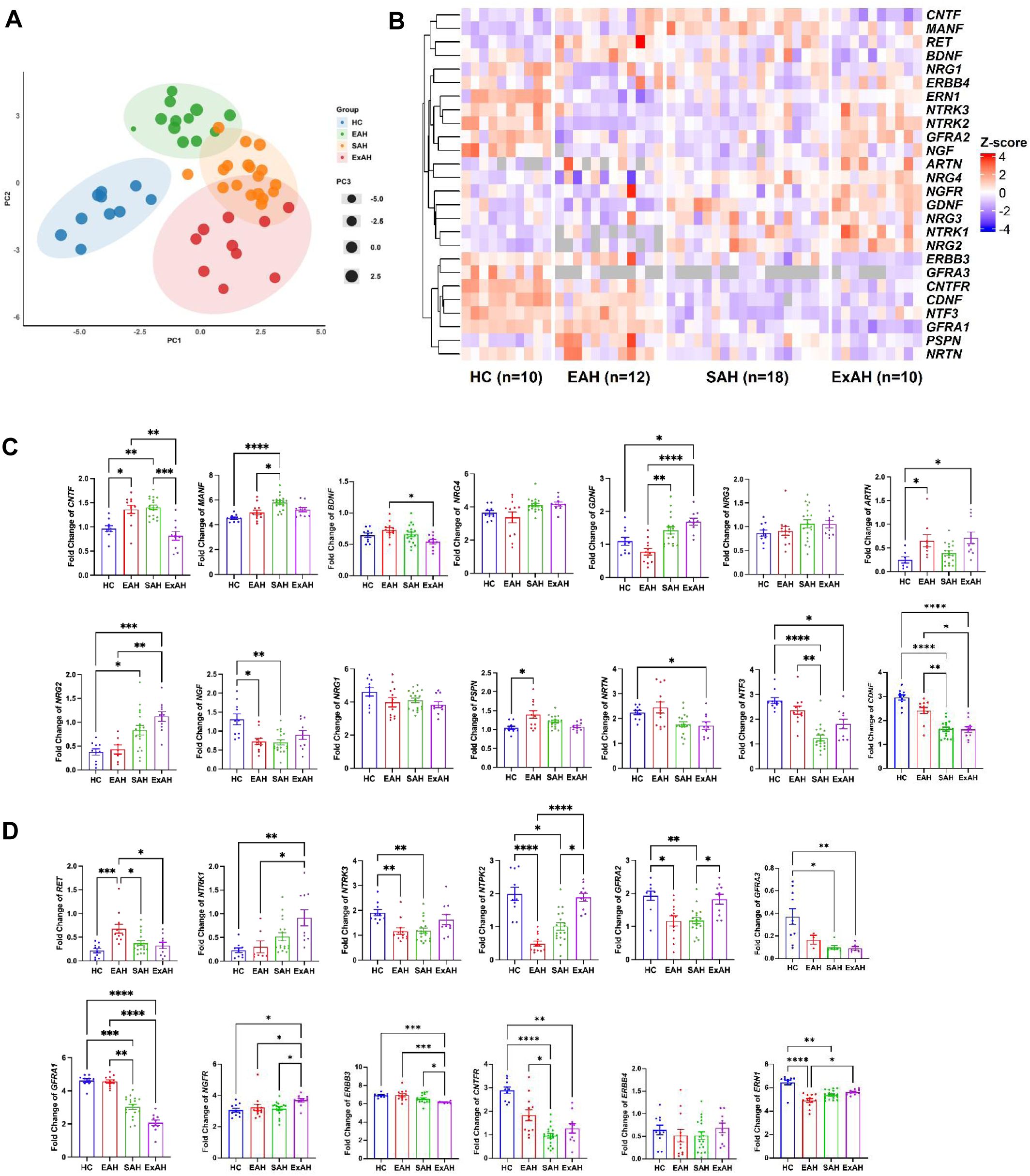
Stage-specific transcriptional remodeling of neurotrophic signaling in human alcoholic liver disease. (**A**) Principal component analysis (PCA) of 26 neurotrophic factor–related genes across healthy controls (HC, n=18), early alcoholic hepatitis (EAH, n=12), severe alcoholic hepatitis (SAH, n=18), and explanted alcoholic livers (ExAH, n=10). (**B**) Heatmap of Z-score normalized expression of neurotrophic factors and their receptors across ALD stages. (**C**) Expression profiles of neurotrophic factors across disease stages. (**D**) Expression profiles of neurotrophic factor receptors across disease stages. Data is derived from a publicly available human liver bulk RNA-seq dataset. Each dot represents an individual sample. Statistical analysis was performed using Kruskal–Wallis test followed by Dunn’s multiple-comparison test. *\*P < 0.05, **P < 0.01, ***P < 0.001, ****P < 0.0001*.

### Neurotrophic factors are associated with clinical severity and inflammatory signatures in patients with ALD

To further determine whether dysregulated neurotrophic signaling is linked to injury outcomes and immune responses in patients with ALD, we performed correlation analyses between neurotrophic genes and key clinical parameters as well as inflammation– and fibrosis-related genes. As shown in **Figure 3A**, several neurotrophic factors, including *MANF*, *GDNF*, *NTRK1*, *NRG2*, and *NRG4*, were positively correlated with multiple clinical indicators of disease severity such as Model for End-stage Liver Disease (MELD) score, the Age, serum Bilirubin, INR, and serum Creatinine (ABIC) score, total bilirubin (TBIL), alkaline phosphatase (ALP) and international normalized ratio (INR). Meanwhile, these factors exhibited negative correlations with clinical markers of hepatic function and recovery, such as platelet count and albumin levels. Conversely, factors, such as *NRTN*, *GFRA3*, *NTF3*, *CDNF* and *GFRA1*, displayed the opposite pattern, showing negative correlations with severity markers but positive correlations with protective indicators. Some factors including *NTRK2*, *NGF*, *NTRK3* and *GFRA2* were positively associated only with serum injury markers such as AST and ALT.

**Figure 3.**
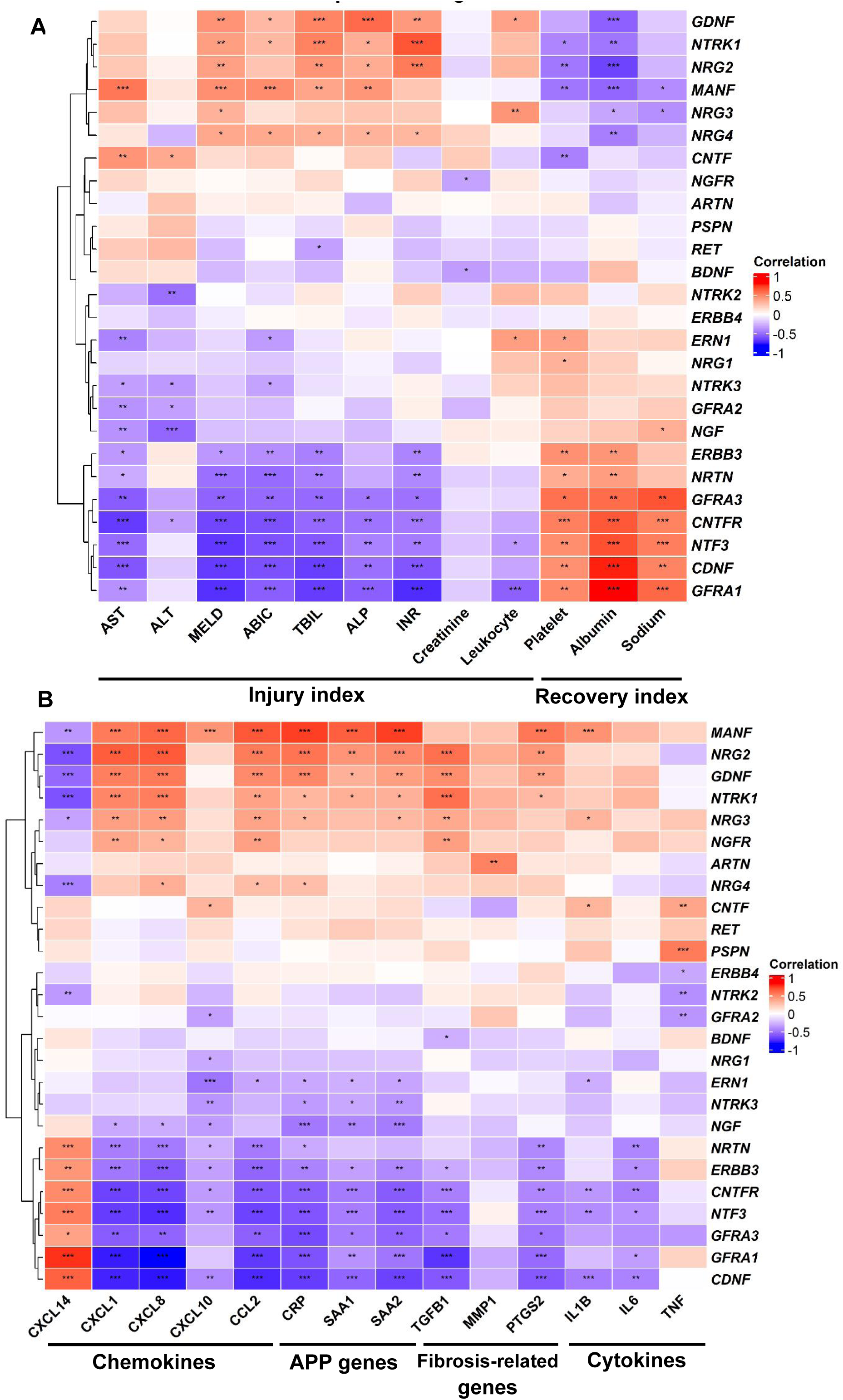
Neurotrophic factors are associated with clinical severity and inflammatory signatures in human alcoholic liver disease. (**A**) Heatmap showing correlations between neurotrophic factor–related genes and clinical parameters across ALD patients, including markers of liver injury, disease severity, and recovery. (**B**) Heatmap showing correlations between neurotrophic factor–related genes and inflammatory and fibrosis-related gene expression, including chemokines, acute-phase proteins, fibrosis-related genes, and cytokines. Correlations were calculated using Spearman’s rank correlation. Statistical significance is indicated as follows: *\*P < 0.05, **P < 0.01, ***P < 0.001*.

More interestingly, most neurotrophic factors were also strongly correlated with pro-inflammatory and fibrosis-related genes (**Figure 3B**). For example, the positively injury-correlated factors including *MANF*, *NRG2*, *GDNF*, *NTRK1* and *NRG3* showed positive correlations with multiple chemokines (e.g. *CXCL1*, *CXCL8*, *CCL2*), acute-phase proteins (APPs) (e.g. *CRP*, *SAA1, SAA2*), and fibrosis-related genes (e.g. *TGFβ1*, *MMP9, PTGS2*). In contrast, negatively injury-correlated factors including *NGF*, *NRTN*, *GFRA3*, *GFRA1*, *NTF3* and *CDNF* were inversely associated with these pro-inflammatory and fibrotic markers. Consistently, these dysregulated factors exhibited the opposite trend with an anti-inflammatory chemokine *CXCL14* (14,15).

In summary, these data reveal that distinct neurotrophic factors are tightly linked to both clinical severity and inflammatory signaling in ALD, underscoring their potential roles as modulators of disease progression.

### Co-expression clustering and stage-dependent rewiring of neurotrophic signaling during ALD progression

While differential expression analyses reveal which neurotrophic genes change with disease progression, they do not capture how these genes interact or whether they operate within coordinated signaling circuits. To better understand whether neurotrophic factors exhibit coordinated expression and potential functional relatedness, we performed pairwise gene-gene correlation analysis among these 26 neurotrophic factors using Pearson correlation. The heatmap revealed a pronounced modular organization rather than a single, uniform co-expression network (**Figure 4A**). Hierarchical clustering identified multiple relatively coherent gene modules with distinct correlation structures. One prominent co-expression module, located in the upper-left region of the heatmap, comprised *NTRK3, ERN1, NGF, NTRK2, GFRA2, NRG1* and *ERBB4*, which exhibited strong positive correlations with one another, forming a tightly associated subset. A second module, centered in the middle of the heatmap, included *CNTFR, NTF3, CDNF, GFRA3*, *NRTN, ERBB3* and *GFRA1*, displaying coordinated expression across samples and constituting a distinct correlation block. In contrast, a smaller module in the lower-right region, represented by *MANF, NRG4, GDNF* and *NRG2*, showed positive correlations within the group but predominantly weak or negative correlations with genes from the other clusters, indicating a transcriptionally distinct subset. Together, these findings highlight a modular architecture of neurotrophic gene expression, consistent with divergent regulatory programs and potentially distinct functional roles during ALD progression.

**Figure 4.**
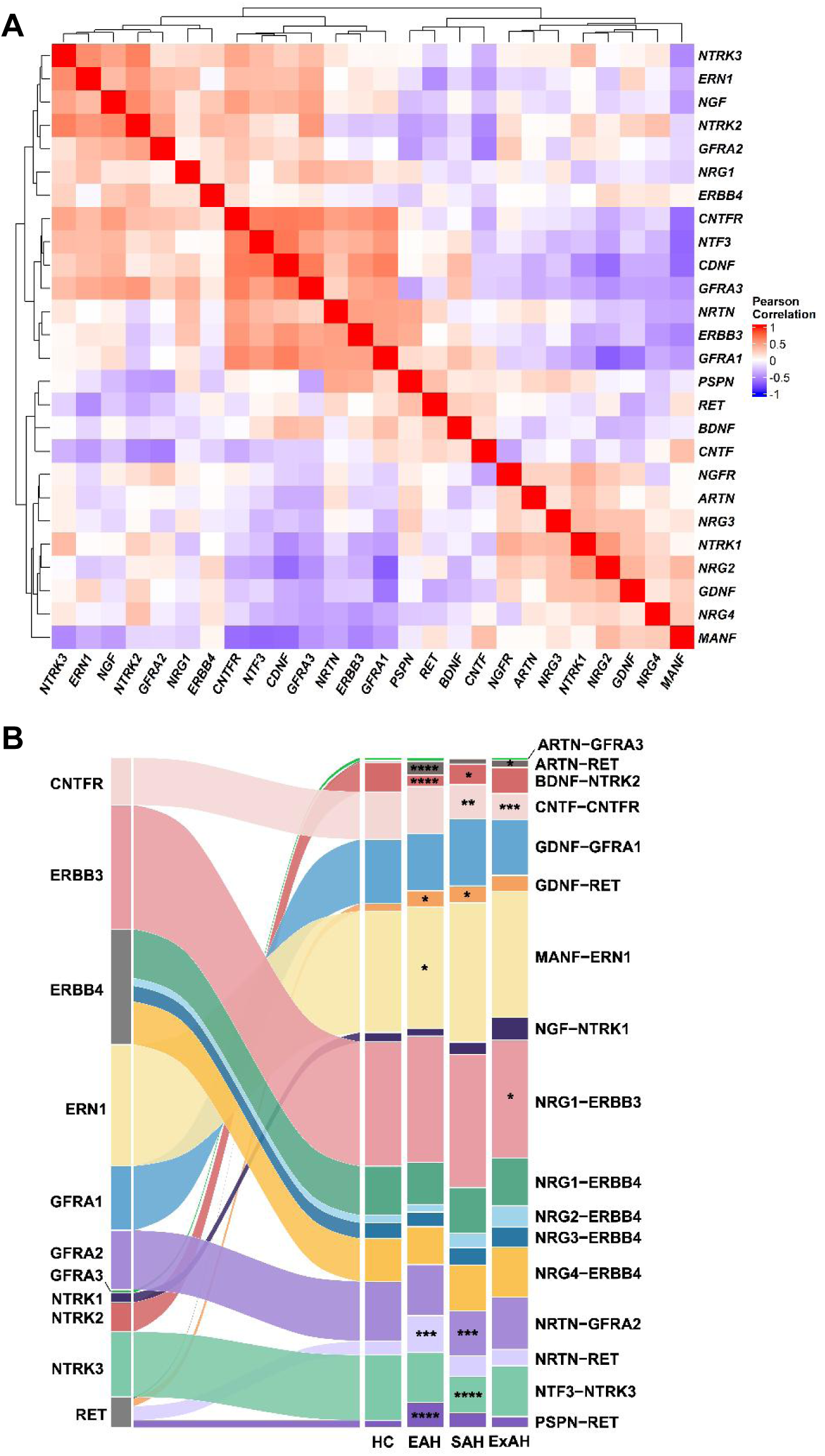
Co-expression clustering and ligand–receptor relationships of neurotrophic factors during ALD progression. (**A**) Heatmap showing pairwise Pearson correlations among neurotrophic factor–related genes with hierarchical clustering. (**B**) Alluvial plot showing the relative contribution of neurotrophic ligand–receptor pairs across ALD stages (HC, EAH, SAH, ExAH). Flow width represents the proportional contribution of each ligand– receptor pair in each stage. Correlation coefficients are indicated by the color scale. Statistical significance was determined using the Kruskal–Wallis test followed by Dunn’s multiple-comparison test with Benjamini–Hochberg correction. *\*P < 0.05, **P < 0.01, ***P < 0.001*.

To further assess how neurotrophic ligand-receptor relationships evolve across disease stages, we visualized the relative contribution of individual ligand-receptor pairs using an alluvial plot, which can reveal dynamic redistribution and selective rewiring of interactions during ALD progression (**Figure 4B**). Several ligand-receptor pairs maintained relatively stable and prominent contributions throughout disease progression. Notably, the MANF-ERN1 and NRG1-ERBB3 axes consistently accounted for a substantial proportion of the overall signal across all stages. In contrast, other ligand-receptor pairs exhibited progressive, stage-associated changes. The contribution of NRG2-ERBB4 increased across successive stages, accompanied by a gradual expansion of NRTN-GFRA2, indicating a stage-associated increase in their relative representation. Additional pairs demonstrated gradual shifts in contribution with disease progression; for example, CNTF-CNTFR decreased at later stages, whereas GDNF-RET showed a consistent increase in AH compared to HCs. Beyond monotonic trends, several ligand-receptor pairs displayed non-linear dynamics. NRTN-RET and PSPN-RET exhibited increased representation at intermediate stages followed by a decline at later stages, whereas BDNF-NTRK2 and NGF-NTRK1 showed an initial decrease at intermediate stages with partial recovery at later stages. Collectively, these findings indicate that neurotrophic ligand-receptor interactions undergo dynamic, pair-specific reorganization across ALD stages.

### Cell-specific dysregulation of neurotrophic factors in patients with AH

To identify the cellular sources underlying altered neurotrophic signaling in AH, we integrated and analyzed publicly available single-cell RNA sequencing (scRNA-seq) datasets (GSE136103 & GSE255772) (16, 17) comprising hepatic non-parenchymal cells (NPCs) from 5 HCs and 5 patients with AH. After quality control, a total of 51,273 cells were included for downstream analysis. To define the cellular landscape, we performed unsupervised clustering followed by annotation based on canonical marker gene expression (**supplemental Figure S1A**). Uniform manifold approximation and projection (UMAP) revealed distinct clusters corresponding to major hepatic cell populations, including immune cells, liver sinusoidal endothelial cells (LSECs) and hepatic stellate cells (HSCs), along with a limited number of hepatocytes (**Figure 5A**).

**Figure 5.**
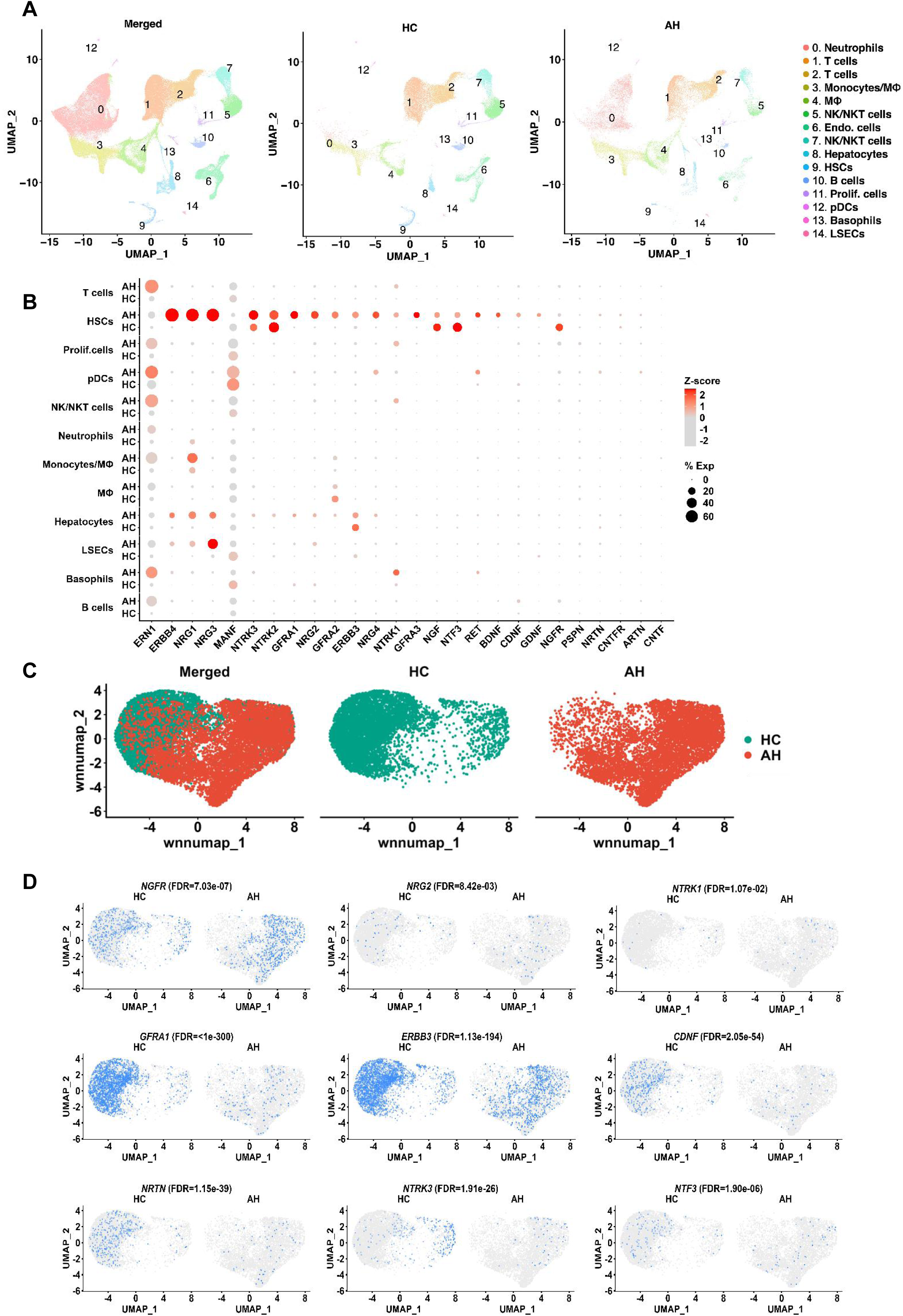
Cell type-specific expression of neurotrophic factors in patients with AH. (**A**) UMAP plots showing the distribution of major hepatic cell populations across all cells (Merged), HC, and AH samples, with clusters annotated based on canonical marker gene expression. (**B**) Dot plot showing the expression of neurotrophic factor–related genes across annotated cell types in HC and AH samples. Dot size represents the percentage of expressing cells, and color indicates scaled expression levels. (**C**) WNN-UMAP plots showing the distribution of hepatocyte-enriched spots in merged samples (Merged), HC, and AH samples from the human spatial transcriptomic dataset. (**D**) WNN-UMAP FeaturePlot visualization of neurotrophic factor–related genes in hepatocyte-enriched spots from the human spatial transcriptomic dataset. Blue dots indicate gene-positive spots and gray dots indicate non-expressing spots. Differential expression between HC and AH groups was assessed using the Wilcoxon rank-sum test followed by Benjamini–Hochberg FDR correction.

We next sought to delineate cell-specific alterations in neurotrophic signaling by performing differential expression analysis between AH and HC samples within each annotated population. Dotplot visualization (**Figure 5B**) demonstrated that the majority of trophic genes were enriched in healthy HSCs, which was further enhanced in AH HSCs. Notably, *ERN1* and *MANF* exhibited broad expression across multiple cell types, suggesting a potentially conserved role in hepatic stress responses. In contrast, several neurotrophic ligands and receptors displayed selective enrichment within stromal and immune compartments. For example, *NRG1* was predominantly expressed in monocytes/macrophages and further elevated in AH, whereas *NRG3* was highly induced in AH endothelial cells. These findings indicate specialized contributions of NPC population to neurotrophic signaling in the AH liver.

Since hepatocytes were underrepresented in the scRNA-seq datasets, we further evaluated neurotrophic gene expression using an independent human liver spatial transcriptomic dataset (GSE278662) containing hepatocyte-enriched spots from both HC (donors n=2, spots n= 6651) and AH (patients n=2, spots n=5993) samples **(Figure 5C)** (18). UMAP analysis of hepatocyte-enriched spots revealed heterogeneous and gene-specific expression patterns, highlighting a distinct neurotrophic landscape within parenchymal cells **(Figure 5D, Supplemental Table S2)**. Among the most significantly altered markers, the neurotrophic receptors NGFR and NTRK1, as well as the ligand NRG2, were markedly upregulated in AH hepatocytes. In contrast, expression of the neurotrophic receptors GFRA1, ERBB3 and NTRK3, as well as the ligands CDNF, NRTN and NTF3, was significantly reduced in AH hepatocytes. These findings were consistent with observation from the Ramon dataset.

Collectively, these results demonstrate that neurotrophic signaling in the human liver is highly cell-specific and undergoes selective remodeling in AH, with prominent contributions from HSCs and other non-parenchymal compartments. These findings provide a cellular framework for understanding how neurotrophic pathways may coordinate intercellular communication and stress adaptation during ALD progression.

### Impaired hepatic NTF3 signaling and sympathetic innervation in human AH

To validate our transcriptomic findings, we examined human liver tissues from an independent cohort of HCs and patients with AH. Clinicopathological characteristics of the study subjects are summarized in **supplemental Table S1**. At the protein level, Western blot analysis further validated that parasympathetic marker ChAT and sympathetic marker TH were both decreased in AH, while the pan-neuron marker PGP9.5/UCHL1 was elevated in AH (**Figure 6**), supporting the transcriptomic indication of neural remodeling.

**Figure 6.**
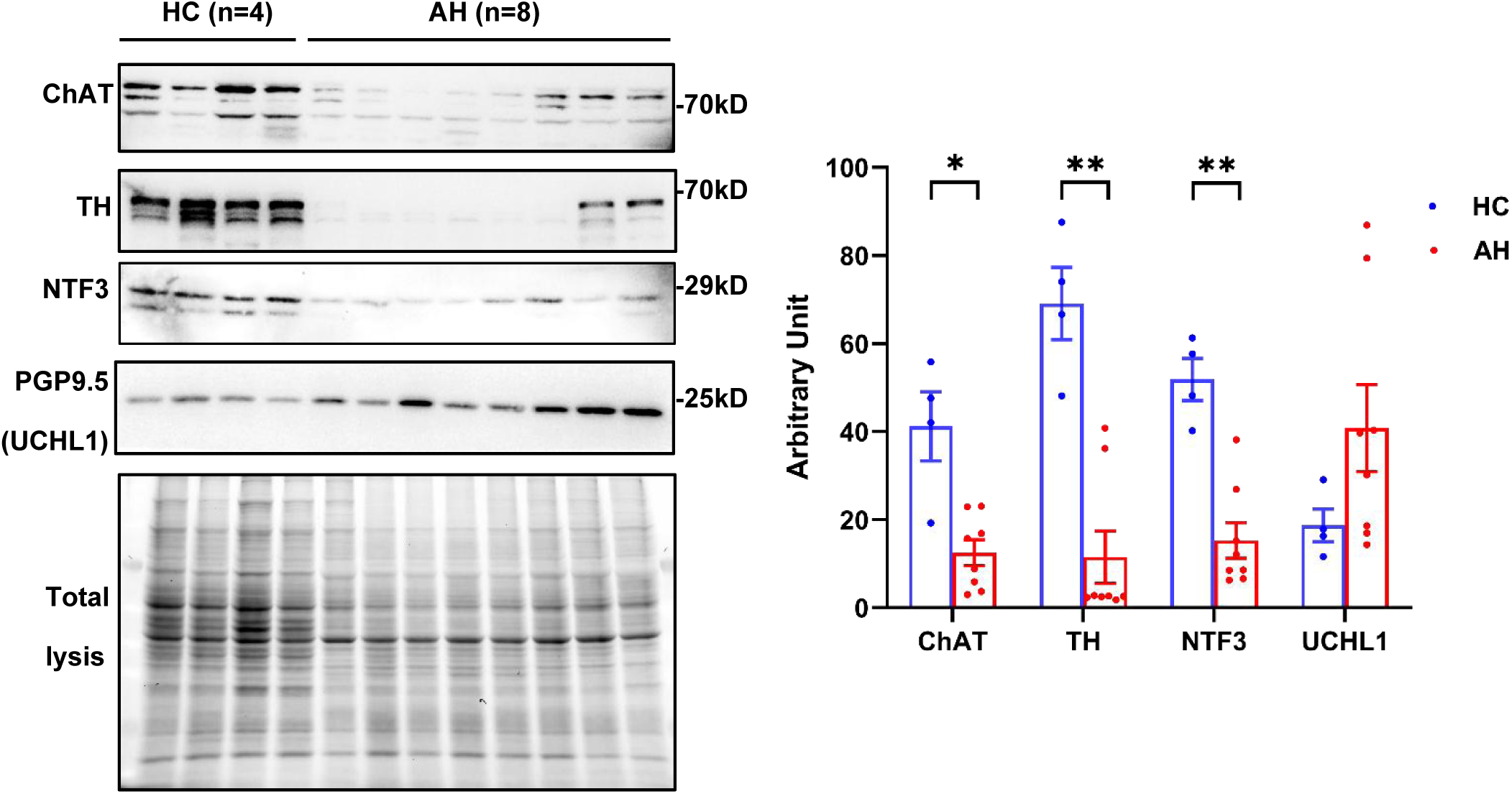
Validation of dysregulated neurotrophic and neural programs in human AH. Representative Western blot images and quantification of neural markers in HC (n=4) and AH (n=8). Data are presented as mean ± SEM. Statistical analysis was performed using the Mann–Whitney U test. *\*P < 0.05, **P < 0.01*.

Given the importance of neurotrophic signaling in maintaining neural integrity, we further assessed the expression of NTF3, a key regulator of sympathetic neuron survival and axonal growth (19). Notably, NTF3 protein levels were markedly reduced in AH compared with HCs. This finding suggests that chronic ethanol exposure may impair hepatic NTF3-mediated neurotrophic support, thereby contributing to the disruption of sympathetic innervation observed in AH.

Collectively, these results validate the computational predictions from bulk and single-cell transcriptomic analyses and demonstrate that neurotrophic and neural gene programs are coordinately modulated in human ALD. The concurrent upregulation of multiple neurotrophic factors alongside neural markers suggests a dynamic remodeling of hepatic neural networks in response to alcohol-induced injury. These findings reinforce the concept that neurotrophic signaling is intimately linked to hepatic stress responses and may represent a previously underappreciated mechanism connecting neural plasticity with liver pathology in ALD.

## Discussion

Emerging evidence indicates that neurotrophic factors have important extra-neural roles in peripheral tissues, including the liver, where they contribute to energy homeostasis and insulin sensitivity (20, 21). Hepatic function is tightly regulated by both branches of the autonomic nervous system through aminergic, peptidergic, adrenergic and cholinergic innervation (20, 22, 23). Sympathetic fibers primarily supply nerve endings along hepatic lobules and sinusoids, whereas parasympathetic (cholinergic) fibers penetrate deeper into the hepatic parenchyma in both humans and mice (24–27). Despite this intricate neural regulation, few studies have examined how autonomic innervation and neurotrophic signaling are altered during ALD progression. In this study, we demonstrate that autonomic nerve and neurotrophic signaling undergo stage-dependent transcriptional remodeling across the spectrum of human ALD. This reprogramming is closely linked to clinical indicators of disease severity, inflammatory signatures and fibrotic remodeling. Importantly, our findings highlight that neurotrophic pathways and neural networks are not merely bystanders but integral components of the ALD pathogenic landscape. By uncovering these coordinated changes, we expand the current conceptual framework of alcohol-induced liver injury to include a neuro-regulatory dimension, suggesting that hepatic neural remodeling may actively contribute to disease progression and represent a novel therapeutic axis.

Neurotrophic factors have been increasingly implicated in context-dependent metabolic adaptation. For example, GDNF, one of most extensively studied trophic factor (28), is elevated in livers of patients with advanced fibrosis associated with metabolic dysfunction-associated steatohepatitis (MASH; 27), ALD (29) and viral hepatitis (30) as well as in multiple murine models of liver injury, including high fat diet (HFD) feeding (31), methionine-choline deficient (MCD) diet feeding (29), bile duct ligation (BDL; 29) or carbon tetrachloride (CCl4) treatment (29). Notably, GDNF appears to exert context-dependent effects: it protects against HFD-induced obesity, hepatic steatosis, inflammation, and fibrogenesis (31), yet promotes pro-inflammatory gene expression and exacerbates fibrosis in BDL and CCl₄ models (29). Consistent with this dichotomy, we observed a progressive increase in GDNF expression across ALD stages, accompanied by positive correlations with clinical severity as well as inflammatory and fibrotic signatures. These findings suggest that GDNF may play a pathogenic, rather than protective, role in ethanol-induced liver injury. Similarly, *NRG2* expression increased steadily with ALD progression. While neuregulin family members such as NRG4 are well recognized for their roles in fatty liver disease (32, 33) and NRG1 has been implicated in triglyceride regulation during steatosis (24,35), the specific function of NRG2 in liver disease, including ALD, remains poorly defined, compared to its established roles in the nervous system. In contrast, *CDTF*, an unconventional trophic factor and negative regulator of the unfolded protein response (UPR) with anti-inflammatory (36–38), exhibited a decline during ALD progression, consistent with loss of protective signaling (37). Interestingly, *CNTF* was upregulated as early as EAH, suggesting an early stress-responsive activation, whereas induction of *NTRK1* was observed predominantly at the explanted end stage, potentially reflecting a late decompensatory response. Together, these findings indicate that ALD is characterized not only by activation of stress-associated trophic pathways but also by stage-specific reprogramming of neurotrophic signaling networks induced by chronic alcohol exposure.

Autonomic imbalance is increasingly recognized as a shared feature across chronic liver diseases, including ALD. Neurotrophic signaling is essential for maintaining neural integrity in peripheral tissues, and our data point to a potential role for NTF3 in preserving hepatic sympathetic innervation. In particular, we observed a coordinated downregulation of NTF3 and canonical sympathetic nerve markers in AH, suggesting impaired neurotrophic support contributes to sympathetic dysfunction. Accumulating evidence suggests that metabolic stress disrupts hepatic neural integrity, leading to sympathetic nerve degeneration and architectural disorganization (7, 8). Interestingly, this process appears to be driven, at least in part, by neuro-immune crosstalk, as macrophage-derived TNF-α has been shown to contribute to sympathetic nerve injury (7). A similar mechanism is likely operative in the context of ethanol-induced liver injury. Consistent with this notion, we observed that canonical sympathetic nerve markers, including DBH and TH, were significantly downregulated in patients with AH in two independent cohorts, indicating impaired sympathetic signaling. Notably, the expression of these markers was inversely correlated with TNF-α levels. Given that ethanol exposure robustly induces TNF-α production in both human and experimental models (39–41), further supporting a role for inflammatory mediators in driving neural dysfunction. However, sympathetic hyperactivity has been reported in advanced liver disease, including ALD (42, 43). In line with this, chronic ethanol feeding in mice increases the levels of catecholamines, such as epinephrine and norepinephrine, in the cecum, portal circulation and liver. This paradoxical elevation of neurotransmitters, despite reduced neural markers, suggests dysregulated or ectopic catecholamine production, which has been linked to increased apoptosis of Kupffer cells (44). Taken together, these findings support a model in which impaired NTF3-mediated neurotrophic support and inflammation-driven neural injury converge to disrupt hepatic sympathetic integrity in ALD. Concurrently, altered catecholamine dynamics may further modulate immune cell survival and function. This bidirectional neuro-immune interplay represents a key mechanism linking metabolic stress, inflammation, and tissue injury during chronic alcohol exposure.

HSCs are increasingly recognized as functionally analogous to neural pericytes or astrocytes and are often referred to as “hepatic neuroglia” due to their expression of neural-associated markers (45–47). Consistently, our scRNA-seq analysis of human NPCs demonstrated that a substantial proportion of neurotrophic genes are enriched in HSCs in both HCs and AH. Notably, ethanol-induced activation of HSCs further enhances their capacity to produce neurotrophic factors, suggesting that these cells may serve as a major source of neurotrophic signaling in the injured liver. In turn, neurotrophic factors can modulate HSC activation and fibrogenic responses, although their effects appear context-dependent, with evidence supporting both protective and profibrotic roles (29, 31, 48). In parallel, sympathetic overactivity has been shown to promote HSC proliferation and collagen production through catecholamine-mediated signalin (49, 50). Together, these findings position HSCs at the center of a dynamic neuro-stromal axis in ALD, where neurotrophic signaling and autonomic inputs converge to regulate stellate cell activation and fibrosis progression.

Several limitations should be acknowledged. First, the present study is primarily based on transcriptomic analyses and correlation-based inference, which precludes definitive conclusions regarding causality. Although strong associations were observed between neurotrophic factors, inflammatory signatures, and clinical severity, functional validation will be required to determine whether these pathways actively drive disease progression or represent secondary responses to tissue injury. Second, bulk RNA-seq lacks spatial resolution, and while single-cell analysis provides cellular context, it does not capture the anatomical organization of intrahepatic neural circuits. Spatial transcriptomics and imaging-based approaches would be valuable for delineating the precise localization and structural remodeling of neural elements during ALD progression. Third, although the data span multiple disease stages, longitudinal sampling within the same individuals was not available, limiting inference about temporal dynamics. Finally, although the present study is based on human datasets and therefore directly relevant to clinical disease, mechanistic validation will be required to determine how specific neurotrophic pathways functionally influence hepatic inflammation and injury. Experimental perturbation studies, whether in human-relevant model systems or complementary animal models, will be necessary to establish causal relationships and evaluate therapeutic potential.

In summary, our study demonstrates that ALD is accompanied by structured, stage-dependent remodeling of neurotrophic and autonomic signaling networks that are tightly integrated with inflammatory amplification and clinical severity. Rather than reflecting a uniform response, ALD exhibits a pattern of bidirectional neurotrophic divergence, modular network reorganization, and coordinated dysregulation of neural markers. These findings position trophic pathways as integral components of the neuro-immune landscape in ethanol-induced liver injury and suggest that disruption of neurotrophic-autonomic homeostasis may represent a previously underappreciated dimension of ALD pathophysiology. By linking neural remodeling to inflammation and disease progression, our work provides a conceptual framework for understanding how neuro-regulatory mechanisms contribute to ethanol-induced liver injury and highlights potential avenues for therapeutic intervention.

## Methods

### Human liver samples

Liver tissue samples from patients with severe alcohol-associated hepatitis (AH, n = 8) undergoing liver transplantation and wedge biopsies from healthy donor livers (HC, n = 3–4) were collected, snap-frozen in liquid nitrogen, and stored at −80°C. Samples were obtained from the NIAAA R24 Clinical Resource for Alcoholic Hepatitis Investigations at Johns Hopkins University. Clinical information is provided in **Supplemental Table S1**. Both male and female patients were included in the study. Analyses were performed without sex-based exclusion.

### Human liver RNA-seq data acquisition and preprocessing

Publicly available bulk RNA-seq data of human liver tissues were obtained from the Pitt–Ramon cohort (51), including healthy controls (HC, n = 10), early alcohol-associated hepatitis (EAH, n = 12), severe alcohol-associated hepatitis (SAH, n = 18), and explanted end-stage alcohol-associated hepatitis (ExAH, n = 10). Sex was not used as a selection criterion. Raw read counts were normalized to transcripts per million (TPM) and log₂ –transformed. Principal component analysis (PCA) and hierarchical clustering were performed using the *prcomp* function in R (version 4.3.2).

### Analysis of Neurotrophic Factor Expression Signatures

A panel of 26 neurotrophic factor genes and receptors (e.g., MANF, GDNF, NRG2, NTRK1-3, GFRA1-3, NTF3, CNTF, CDNF, and ARTN) was used for expression analysis. Heatmaps were generated using the *ComplexHeatmap* package (version 2.20.0) in R, with z-score normalization applied across genes. Genes with zero expression or missing values across all samples were excluded prior to visualization. Differences among groups (HC, EAH, SAH, and ExAH) were assessed using the Kruskal-Wallis test followed by Dunn’s multiple-comparison correction in GraphPad Prism (version 10.2). Adjusted *P* values < 0.05 were considered statistically significant.

### Correlation analysis with clinical parameters and chemokine genes

Correlations between neurotrophic factor expression and clinical variables were assessed using the *psych* package (Pearson’s method) in R. Clinical indices included AST, ALT, MELD, ABIC, TBIL, ALP, INR, creatinine, leukocyte and platelet counts, albumin, sodium, age, and sex. Correlation coefficients were visualized as heatmaps using *ComplexHeatmap* and *circlize*. Statistical significance was annotated directly on the heatmaps. Correlation analyses with chemokine genes were performed using the same statistical and visualization workflow. The analysis of neural and neuroplasticity markers (e.g., *TH, DBH, TUBB3, STMN2, UCHL1,* and *S100B*) followed the same workflow as for neurotrophic factors, including z-score normalization, statistical comparison among disease stages, and correlation with clinical parameters.

### Co-expression network and ligand–receptor interaction analysis

Correlations between neurotrophic factor expression and clinical variables were assessed using Pearson’s method implemented in the psych package in R. Clinical variables included AST, ALT, MELD, ABIC, TBIL, ALP, INR, creatinine, leukocyte and platelet counts, albumin, sodium, age, and sex. Correlation coefficients were visualized using *ComplexHeatma*p and *circlize,*with statistical significance annotated. Correlation analyses with chemokine genes were performed using the same approach. Analyses of neural and neuroplasticity markers (e.g., TH, DBH, TUBB3, STMN2, UCHL1, and S100B) were conducted using the same workflow.

### Single-cell RNA-seq analysis

Single-cell RNA-seq analysis was performed using publicly available datasets from GSE136103 and GSE255772. Further analyses were conducted using the Seurat package (version 5.4.0). Cells with a high proportion of mitochondrial gene expression (>10%), as well as low-quality cells with fewer than 200 or more than 7,500 detected features, were excluded from downstream analyses. Filtered cells were normalized, and highly variable genes were identified using the FindVariableFeatures function. Data integration was subsequently performed using the reciprocal principal component analysis (RPCA) workflow implemented in Seurat through the IntegrateData function. For clustering analysis, the first 30 principal components (PCs) and a clustering resolution of 0.12 were used. Cell clustering was performed using the FindNeighbors and FindClusters functions. Differentially expressed genes were identified using the FindMarkers function. A total of 51,273 cells derived from HC (n = 5) in GSE136103 and AH (n = 5) samples in GSE255772 were included for downstream analyses. Finally, UMAP plots, dot plots, feature plots, and heatmaps were generated in R.

### Spatial transcriptomic

Spatial transcriptomic data from AH and non-ALD control liver tissues were obtained from the publicly available dataset GSE278662. AH patients (n=2) and non-ALD controls (donors) (n=2) were used for Visium spatial transcriptomic. Previously published spatial spot annotations and cell type classifications from the original study were retained for downstream analyses. Weighted nearest neighbor (WNN)-UMAP analysis integrating RNA and protein modalities was used for dimensionality reduction and visualization. Neurotrophic factor–related genes were analyzed using FeaturePlot– and UMAP-based approaches. Differential expression between HC and AH groups was assessed using the Wilcoxon rank-sum test with Benjamini–Hochberg false discovery rate (FDR) correction. Plots were generated in R.

### Western blot analysis

Protein expression was assessed by Western blot using total protein extracted from frozen human liver tissues from HC and AH patients. Protein concentrations were determined using a BCA Protein Assay Kit (Thermo Fisher Scientific). Equal amounts of protein were resolved by SDS–PAGE (10%) and transferred onto PVDF membranes (Millipore). Membranes were blocked with 5% nonfat milk and then incubated overnight at 4°C with primary antibodies, including rabbit anti-TH (1:1000, AB152, Abcam), rabbit anti-CHAT (1:1000, A13244, ABclonal), and rabbit anti-PGP9.5/UCHL1 (1:20000, A19101, ABclonal), NTF3 (1:500, A12476, ABclonal). followed by incubation with HRP-conjugated anti-rabbit IgG (1:5000, AS014, ABclonal). Protein signals were detected using an enhanced chemiluminescence reagent (Thermo Fisher Scientific) and imaged using a ChemiDoc MP Imaging System (Bio-Rad). Densitometric quantification was normalized to total protein staining using trichloroethanol (TCE)-based labeling to correct for loading differences.

### Statistics

Statistical analyses were performed using R (version 4.3.2) and GraphPad Prism (version 10.2). Specific statistical tests used for each analysis are described in the corresponding figure legends and Methods sections. Data are presented as mean ± SEM unless otherwise indicated. Comparisons between two groups were performed using the Mann–Whitney U test, while comparisons among multiple groups were conducted using the Kruskal–Wallis test followed by Dunn’s multiple-comparison correction. Correlation analyses were performed using Pearson’s correlation coefficient. A two-tailed *P* value < 0.05 was considered statistically significant.

## Author contributions

YZ and XW contributed to the conception and design of the manuscript. YZ, JL, FZ and MSD contributed to the collection and assembly of the data. YJ, YG, TW, YM, SM CT, BG and XW contributed to data analysis and interpretation. YZ, GD, MH and XW contributed to the manuscript writing. All authors gave final approval of the manuscript.

## Acknowledgements

This work was supported by the National Institutes of Health (NIH) grant R00 AA029146 (XW), the Texas Digestive Disease Center (DDC) grant P30 DK056338 (XW), the American Gastroenterological Association (AGA) Pilot Research Award AGA2025-21-06 (XW) and the startup funds from the University of Texas Health Science Center at Houston. We thank the NIAAA Clinical Resources for AH Investigators (R24AA025017, Z.S.) at Johns Hopkins for providing the human samples used in this study.

## References

1. Devarbhavi H, et al. Global burden of liver disease: 2023 update. J Hepatol. 2023;79(2):516–537.

2. Danpanichkul P, et al. Global epidemiology of alcohol-related liver disease, liver cancer, and alcohol use disorder, 2000–2021. Clin Mol Hepatol. 2025;31(2):525–547.

3. Wangler S, et al. The autonomic nervous system in the regulation of glucose and lipid metabolism. Nat Rev Endocrinol. 2026;22:274–286.

4. Spaak J, et al. Dose-related effects of red wine and alcohol on heart rate variability. Am J Physiol Heart Circ Physiol. 2010 Jun;298(6):H2226–31.

5. Wickramasinghe PB, et al. Ablation of sympathetic nerve–β3 adrenergic receptor– mediated adipose tissue lipolysis attenuates alcohol-induced liver injury in mice. Cell Mol Gastroenterol Hepatol. 2026;20(1):101623.

6. Song Q, et al. mTORC1 inhibition uncouples lipolysis and thermogenesis in white adipose tissue to contribute to alcoholic liver disease. Hepatol Commun. 2023;7(3):e0059.

7. Liu K, et al. Metabolic stress drives sympathetic neuropathy within the liver. Cell Metab. 2021;33(3):666–675.e4.

8. Adori C, et al. Disorganization and degeneration of liver sympathetic innervation in nonalcoholic fatty liver disease revealed by 3D imaging. Sci Adv. 2021;7(30):eabg5733.

9. Park H, Poo MM. Neurotrophin regulation of neural circuit development and function. Nat Rev Neurosci. 2013;14(1):7–23.

10. Sousa-Victor P, et al. MANF regulates metabolic and immune homeostasis in ageing and 10. protects against liver damage. Nat Metab. 2019;1(2):276–290.

11. Sopi RB, et al. Role of brain-derived neurotrophic factor in hyperoxia-induced enhancement of contractility and impairment of relaxation in lung parenchyma. Am J Physiol Lung Cell Mol Physiol. 2008;294(2):L348–L355.

12. Zhao P, et al. Gonadotrophin-induced paracrine regulation of human oocyte maturation by BDNF and GDNF secreted by granulosa cells. Hum Reprod. 2011;26(3):695–702.

13. Huang EJ, et al. Neurotrophins: roles in neuronal development and function. Annu Rev Neurosci. 2001;24:677–736.

14. Lv J, et al. CXCL14 overexpression attenuates sepsis-associated acute kidney injury by inhibiting proinflammatory cytokine production. Mediators Inflamm. 2020;2020:2431705.

15. Lee HT, et al. A crucial role of CXCL14 for promoting regulatory T cells activation in stroke. Theranostics. 2017;7(4):855–875.

16. Ramachandran P, et al. Resolving the fibrotic niche of human liver cirrhosis at single-cell level. Nature. 2019;575(7783):512–518.

17. Guan Y, et al. IL-8+ neutrophils drive inexorable inflammation in severe alcohol-associated hepatitis. J Clin Invest. 2024;134(9):e178616.

18. Gripshover TC, et al. Visium spatial transcriptomics and proteomics identifies novel hepatic cell populations and transcriptomic signatures of alcohol-associated hepatitis. Alcohol Clin Exp Res. 2025 Jan;49(1):106–116.

19. Belliveau DJ, et al. NGF and neurotrophin-3 both activate TrkA on sympathetic neurons but differentially regulate survival and neuritogenesis. J Cell Biol. 1997 Jan 27;136(2):375– 388.

20. Amir M, et al. Hepatic autonomic nervous system and neurotrophic factors regulate the pathogenesis and progression of nonalcoholic fatty liver disease. Front Med (Lausanne). 2020;7:62.

21. Iu ECY, et al. Is brain-derived neurotrophic factor a metabolic hormone in peripheral tissues? Biology (Basel). 2022;11(7):1063.

22. Miller BM, et al. Hepatic nervous system in development, regeneration, and disease. Hepatology. 2021;74(6):3513–3522.

23. McCuskey RS. Anatomy of efferent hepatic nerves. Anat Rec A Discov Mol Cell Evol Biol. 2004;280(1):821–826.

24. Das S, et al. The nerve center of organ engineering. Nat Commun. 2025;16(1):9834.

25. Jensen KJ, et al. Hepatic nervous system and neurobiology of the liver. Compr Physiol. 2013;3(2):655–665.

26. Mizuno K, et al. Intrahepatic distribution of nerve fibers and alterations due to fibrosis in diseased liver. PLoS One. 2021;16(4):e0249556.

27. Streba LA, et al. Role of intrahepatic innervation in regulating the activity of liver cells. World J Hepatol. 2014;6(3):137–143.

28. Porcari C, et al. Exploring the diversity of biological processes regulated by glial cell line– derived neurotrophic factor, a pleiotropic molecule with therapeutic potential. Front Physiol. 2025;16:1618330.

29. Tao L, et al. Glial cell line–derived neurotrophic factor mediates hepatic stellate cell activation via ALK5/Smad signaling. Gut. 2019;68(12):2214–2227.

30. Yang G, et al. Serum glial cell line–derived neurotrophic factor is a novel biomarker predicting cirrhosis in chronic hepatitis B. Can J Gastroenterol Hepatol. 2022;2022:1048104.

31. Mwangi SM, et al. Glial cell line–derived neurotrophic factor protects against high-fat diet–induced hepatic steatosis by suppressing hepatic PPAR-γ expression. Am J Physiol Gastrointest Liver Physiol. 2016;310(2):G103–G116.

32. Guo L, et al. Hepatic neuregulin 4 signaling defines an endocrine checkpoint for steatosis-to-NASH progression. J Clin Invest. 2017;127(12):4449–4461.

33. Tapak MA, et al. The impact of serum neuregulin 4 levels on NAFLD: a systematic review and meta-analysis. BMC Gastroenterol. 2025;25(1):509.

34. Meng D, et al. Roles and mechanisms of NRG1 in modulating NAFLD through ErbB3 signaling. Obes Res Clin Pract. 2021;15(2):145–151.

35. Xu C, et al. Neuregulin1 ameliorates metabolic dysfunction–associated fatty liver disease via ERK/SIRT1 signaling pathways. BMC Gastroenterol. 2025;25(1):47.

36. Graewert MA, et al. Structural basis of CDNF interaction with the UPR regulator GRP78. Nat Commun. 2024;15(1):8175.

37. Arancibia D, et al. CDNF induces the adaptive unfolded protein response and attenuates endoplasmic reticulum stress–induced cell death. Biochim Biophys Acta Mol Cell Res. 2018;1865(11):1579–1589.

38. Jäntti M, et al. Trophic activities of endoplasmic reticulum proteins CDNF and MANF. Cell Tissue Res. 2020;382(1):83–100.

39. Zhou Z, et al. A critical involvement of oxidative stress in acute alcohol-induced hepatic TNF-α production. Am J Pathol. 2003;163(3):1137–1146.

40. Lin HZ, et al. Chronic ethanol consumption induces tumor necrosis factor-α and related cytokines. Alcohol Clin Exp Res. 1998;22(5 Suppl):231S–237S.

41. Wu X, et al. Recent advances in understanding alcohol-associated liver disease. Annu Rev Pathol. 2023;18:411–438.

42. Henriksen JH, et al. The sympathetic nervous system in liver disease. J Hepatol. 1998;29(2):328–341.

43. Willett I, et al. Total and renal sympathetic nervous system activity in alcoholic cirrhosis. J Hepatol. 1985;1(6):639–648.

44. Kim HH, et al. Catecholamine induces Kupffer cell apoptosis via growth differentiation factor 15 in alcohol-associated liver disease. Exp Mol Med. 2023;55(1):158–170.

45. Schachtrup C, et al. Hepatic stellate cells and astrocytes: stars of scar formation and tissue repair. Cell Cycle. 2011;10(11):1764–1771.

46. Sato M, et al. Hepatic stellate cells: unique characteristics in cell biology and phenotype. Cell Struct Funct. 2003;28(2):105–112.

47. Puche JE, et al. Hepatic stellate cells and liver fibrosis. Compr Physiol. 2013;3(4):1473– 1492.

48. Sun TT, et al. Neurotrophic factors stimulate activation of hepatic stellate cells in liver fibrosis. Biochem Biophys Res Commun. 2022;630:167–174.

49. Oben JA, et al. Norepinephrine and neuropeptide Y promote proliferation of hepatic stellate cells. Biochem Biophys Res Commun. 2003;302(4):685–690.

50. Oben JA, et al. Hepatic fibrogenesis requires sympathetic neurotransmitters. Gut. 2004;53(3):438–445.

51. Argemi J, et al. Defective HNF4α-dependent gene expression as a driver of hepatocellular failure in alcoholic hepatitis. Nat Commun. 2019;10:3126.

