## Supplementary Figures and Tables for "Autonomic and neurotrophic signaling in alcohol-associated liver disease"

### Supplemental materials

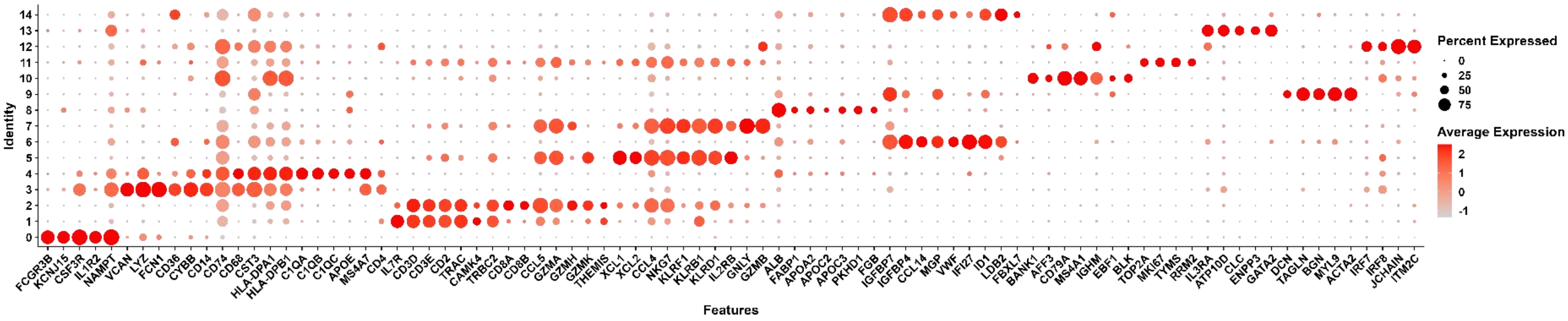


**Figure S1. Cell type annotation of single-cell transcriptomes in human liver.** Unsupervised clustering identified major hepatic cell populations, including neutrophils, monocytes/ macrophages, T cells, NK/NKT cells, hepatocytes, LSECs, B cells, proliferating cells, basophils, HSCs, and pDCs. Cell identities were assigned based on canonical marker gene expression using a curated panel of lineage-specific markers. Dot size represents the percentage of cells expressing each marker gene, and color intensity indicates the scaled average expression level.

**Table S1. Clinicopathological Characteristics of Liver Chunk Samples from Patients with ALD (n=8).**

| Variable | Value |
|---|---|
| Age (years) | 46.00 (38.50–49.00) |
| Male sex, n (%) | 7 (87.5%) |
| Race (White), n (%) | 6 (75.0%) |
| BMI (kg/m²) | 29.66 (26.31–37.01) |
| MELD score | 34.00 (30.00–38.75) |
| INR | 2.19 (1.75–2.87) |
| WBC (×10³/μL) | 14.32 (8.66–21.71) |
| Hemoglobin (g/dL) | 8.95 (8.28–9.27) |
| Platelets (×10³/μL) | 79.50 (65.00–198.25) |
| AST (U/L) | 82.50 (79.50–105.50) |
| ALT (U/L) | 41.50 (28.25–77.25) |
| Total bilirubin (mg/dL) | 28.30 (20.45–44.97) |
| Creatinine (mg/dL) | 2.85 (1.12–3.08) |

The data are presented as median (interquartile range, IQR).

### Supplemental Table S2: Neurotrophic gene expression differences in hepatocyte-enriched spots between HC and AH samples

| Gene | HC_mean | AH_mean | HC_pct | AH_pct | HC_positive_mean | AH_positive_mean | log2FC | p_value | Up_in | FDR | Significance |
|---|---|---|---|---|---|---|---|---|---|---|---|
| *GFRA1* | 0.247372 | 0.038104 | 35.212750 | 2.970132 | 0.702507 | 1.282920 | -2.698653 | 0 | HC | 0 | **** |
| *ERBB3* | 0.351769 | 0.199116 | 46.053225 | 16.152178 | 0.763831 | 1.232749 | -0.821021 | 9.04E-196 | HC | 1.13E-194 | **** |
| *CDNF* | 0.037128 | 0.013814 | 6.705759 | 1.134657 | 0.553667 | 1.217421 | -1.426406 | 2.47E-55 | HC | 2.05E-54 | **** |
| *NRTN* | 0.032387 | 0.013555 | 5.397685 | 1.084599 | 0.600024 | 1.249790 | -1.256585 | 1.84E-40 | HC | 1.15E-39 | **** |
| *GFRA2* | 0.031965 | 0.011452 | 4.796271 | 1.017854 | 0.666457 | 1.125065 | -1.480957 | 6.87E-35 | HC | 3.44E-34 | **** |
| *NTRK3* | 0.033578 | 0.009428 | 3.894151 | 0.901051 | 0.862268 | 1.046382 | -1.832427 | 4.59E-27 | HC | 1.91E-26 | **** |
| *NRG4* | 0.013743 | 0.002319 | 2.255300 | 0.133489 | 0.609363 | 1.737099 | -2.567219 | 9.18E-27 | HC | 3.28E-26 | **** |
| *CNTFR* | 0.023498 | 0.008315 | 3.307773 | 0.700818 | 0.710387 | 1.186504 | -1.498709 | 1.61E-24 | HC | 5.04E-24 | **** |
| *RET* | 0.007608 | 0.002570 | 1.668922 | 0.200234 | 0.455892 | 1.283361 | -1.565999 | 5.11E-17 | HC | 1.42E-16 | **** |
| *NRG1* | 0.069054 | 0.070522 | 10.133815 | 6.157183 | 0.681419 | 1.145368 | 0.030362 | 2.96E-13 | AH | 7.40E-13 | **** |
| *ERBB4* | 0.006947 | 0.000942 | 0.962261 | 0.066745 | 0.721923 | 1.411828 | -2.882061 | 6.43E-12 | HC | 1.46E-11 | **** |
| *MANF* | 0.603013 | 0.601056 | 59.133965 | 40.480561 | 1.019740 | 1.484803 | -0.004688 | 8.56E-12 | HC | 1.78E-11 | **** |
| *NGFR* | 0.059504 | 0.139791 | 9.201624 | 11.213082 | 0.646671 | 1.246681 | 1.232211 | 3.78E-07 | AH | 7.03E-07 | **** |
| *PSPN* | 0.010129 | 0.006112 | 1.443392 | 0.533956 | 0.701749 | 1.144571 | -0.728890 | 3.94E-07 | HC | 7.03E-07 | **** |
| *NTF3* | 0.011950 | 0.010625 | 1.999699 | 0.934423 | 0.597600 | 1.137097 | -0.169532 | 1.14E-06 | HC | 1.90E-06 | **** |
| *NGF* | 0.008864 | 0.005840 | 1.293039 | 0.567329 | 0.685506 | 1.029363 | -0.601999 | 2.91E-05 | HC | 4.55E-05 | **** |
| *NTRK2* | 0.020031 | 0.018203 | 2.405653 | 1.585183 | 0.832672 | 1.148295 | -0.138106 | 0.0012581 | HC | 0.00185 | ** |
| *NRG2* | 0.005915 | 0.012389 | 0.781837 | 1.268146 | 0.756549 | 0.976964 | 1.066653 | 0.0060589 | AH | 0.00842 | ** |
| *NTRK1* | 0.001645 | 0.005185 | 0.255601 | 0.550642 | 0.643688 | 0.941568 | 1.655929 | 0.0081384 | AH | 0.01071 | * |
| *NRG3* | 0.010326 | 0.013027 | 1.714028 | 1.151343 | 0.602412 | 1.131471 | 0.335304 | 0.0099378 | AH | 0.01242 | * |
| *ERN1* | 0.184623 | 0.269754 | 26.883176 | 21.024529 | 0.686759 | 1.283046 | 0.547067 | 0.025712 | AH | 0.03061 | * |
| *ARTN* | 0.000767 | 0.002237 | 0.090212 | 0.200234 | 0.850658 | 1.117178 | 1.543502 | 0.1011132 | AH | 0.1149 | ns |
| *BDNF* | 0.000843 | 0.001915 | 0.105247 | 0.183547 | 0.800810 | 1.043172 | 1.183814 | 0.2434189 | AH | 0.26459 | ns |
| *GDNF* | 0.003101 | 0.004752 | 0.420989 | 0.300350 | 0.736564 | 1.582296 | 0.615999 | 0.265534 | AH | 0.2766 | ns |
| *GFRA3* | 0.000577 | 0.000897 | 0.090212 | 0.083431 | 0.640151 | 1.074774 | 0.634808 | 0.8981932 | AH | 0.89819 | ns |
